# CD300lf conditional knockout mouse reveals strain-specific cellular tropism for murine norovirus

**DOI:** 10.1101/2020.08.19.258467

**Authors:** Vincent R. Graziano, Mia Madel Alfajaro, Cameron O. Schmitz, Renata B. Filler, Madison S. Strine, Jin Wei, Leon L. Hsieh, Megan T. Baldridge, Timothy J. Nice, Sanghyun Lee, Robert C. Orchard, Craig B. Wilen

## Abstract

Noroviruses are a leading cause of gastrointestinal infection in humans and mice. Understanding human norovirus (HuNoV) cell tropism has important implications for our understanding of viral pathogenesis. Murine norovirus (MNoV) is extensively used as a surrogate model for HuNoV. We previously identified CD300lf as the receptor for MNoV. Here, we generated a *Cd300lf* conditional knockout (*CD300lf*^*F/F*^) mouse to elucidate the cell tropism of persistent and non-persistent strains of murine norovirus. Using this mouse model, we demonstrate that CD300lf expression on intestinal epithelial cells (IECs), and on tuft cells in particular, is essential for transmission of the persistent MNoV strain CR6 (MNoV^CR6^) *in vivo*. In contrast, the nonpersistent MNoV strain CW3 (MNoV^CW3^) does not require CD300lf expression on IECs for infection. However, deletion of CD300lf in myelomonocytic cells (*LysM Cre+)* partially reduces CW3 viral load in lymphoid and intestinal tissues. Disruption of CD300lf expression on B cells (*CD19 Cre*), neutrophils (*Mrp8 Cre*), and dendritic cells (*CD11c Cre*) did not affect CW3 viral RNA levels. Finally, we show that the transcription factor STAT1, which is critical for the innate immune response, partially restricts the cell tropism of MNoV^CW3^ to LysM+ cells. Taken together, these data demonstrate that CD300lf expression on tuft cells is essential for MNoV^CR6^, that myelomonocytic cells are a major, but not exclusive, target cell of MNoV^CW3^, and that STAT1 signaling restricts the cellular tropism of MNoV^CW3^. This provides the first genetic system to study the cell type-specific role of CD300lf in norovirus pathogenesis.

**IMPORTANCE:** Human noroviruses (HuNoVs) are a leading cause of gastroenteritis resulting in up to 200,000 deaths each year. The receptor and cell tropism of HuNoV in immunocompetent humans are unclear. We use murine norovirus (MNoV) as a model for HuNoV. We recently identified CD300lf as the sole physiologic receptor for MNoV. Here, we leverage this finding to generate a *Cd300lf* conditional knockout mouse to decipher the contributions of specific cell types to MNoV infection. We demonstrate that persistent MNoV^CR6^ requires CD300lf expression on tuft cells. In contrast, multiple CD300lf+ cell types, dominated by myelomonocytic cells, are sufficient for non-persistent MNoV^CW3^ infection. CD300lf expression on epithelial cells, B cells, neutrophils, and dendritic cells is not critical for MNoV^CW3^ infection. Mortality associated with MNoV^CW3^ strain in *Stat1*^*-/-*^ mice does not require CD300lf expression on LysM+ cells, highlighting that both CD300lf receptor expression and innate immunity regulate MNoV cell tropism *in vivo*.

## INTRODUCTION

Human noroviruses (HuNoV) represent a leading cause of gastroenteritis and an important cause of childhood mortality worldwide (*1, 2*). Noroviruses, a genus within the family *Caliciviridae*, are non-enveloped, positive-sense, single-stranded RNA viruses that are transmitted through the fecal-oral route. Noroviruses are segregated into seven genogroups (GI-GVII) (*3*). Genogroups I, II, and IV contain primarily HuNoVs, while G III, V, VI and VII contain bovine NoVs, murine NoVs (MNoV), feline and canine NoVs (*4*). A complete understanding of the mechanism underlying the pathogenesis and biology of HuNoV infection is still lacking due to limited HuNoV cell culture systems, heterogeneity amongst NoV isolates, and a lack of infectious molecular clones (*4-6*). This has hindered vaccine and antiviral drug development.

Cell tropism is an important determinant of virus transmission, pathogenesis and immune evasion. A detailed molecular understanding of the host and viral determinants underlying norovirus cell tropism are critical for vaccine and therapeutic development (*7*). The cell tropism of noroviruses remains incompletely understood (*8, 9*). HuNoV can replicate in stem cell-derived human intestinal enteroids and human B cell-like cell lines *in vitro* (*10, 11*). Our understanding of HuNoV tropism *in vivo* is largely dependent on samples taken from immunodeficient humans infected with HuNoV or experimentally infected animal models (*9, 12, 13*). HuNoV infection was identified in dendritic cells and B cells of intravenously infected chimpanzees (*5*) while intestinal epithelial cells (IECs), macrophages, lymphocytes, and dendritic cells have been identified as HuNoV target cells in pig models (*14-17*). HuNoV infected intestinal epithelial cells were observed in immunocompromised humans (*18*). More recently, HuNoV was shown to infect enteroendocrine cells, a rare secretory epithelial cell population that plays a critical role in the gut-brain axis (*9*). The determinants of cell tropism including the viral receptor, specific role of bile salts and glycans, remain unclear for HuNoV (*10, 19, 20*).

MNoV represents a model for HuNoV and has enabled identification of important host and viral factors that can regulate NoV replication and pathogenesis *in vitro* and *in vivo* (*12, 21-23*). MNoV can be efficiently propagated *in vitro* and productively infects mice, thus providing a tractable *in vivo* system for NoV studies (*22, 24*). MNoV shares many characteristics with HuNoV, including fecal-oral transmission, capsid structure, intestinal replication, and prolonged shedding after acute infection, reflecting asymptomatic HuNoV infection (*12, 22, 24*). Infectious molecular clones of MNoV have been described with distinct patterns of pathogenesis. MNoV strain CR6 (MNoV^CR6^) causes persistent enteric infection which can spread systemically but fails to induce lethality in type I interferon deficient mice (*25, 26*). In contrast, MNoV strain CW3 (MNoV^CW3^), an infectious molecular clone derived from MNV-1, causes non-persistent systemic infection in immunocompetent mice and lethal infection in mice deficient in type I interferon signaling (*23, 26, 27*).

Though genetically similar, minor genetic variants in these MNoV strains confer distinct *in vivo* phenotypes (*25, 26, 28, 29*). The viral determinants of infection have been mapped to the viral capsid protein VP1 and the viral non-structural protein NS1 (*25-27, 30*). Specific amino acid variants have been identified in VP1 that determine enteric and systemic infection and lethality in mice deficient in type I interferon signaling (*26*). Variants in NS1 enable MNoV^CR6^ to antagonize type III interferon and establish persistent enteric infection (*25, 31*). The host determinants of infection include both the MNoV receptor CD300lf and the innate immune system (*23, 27, 28*). CD300lf is a type I integral membrane protein that binds phospholipids on dead and dying cells and can induce pro- or anti-inflammatory signals depending on the specific context (*32*). CD300lf is expressed on tuft cells and diverse hematopoietic cells including macrophages, dendritic cells, neutrophils, and B cells (*29, 32*). CD300lf has been implicated in the pathogenesis of multiple sclerosis, inflammatory bowel disease, and depression (*33-38*).

The cell tropism of MNoV^CW3^ and MNoV^CR6^ are distinct. Our recent discovery of CD300lf as the receptor for MNoV enabled us to determine that radiation-resistant cells are required for MNoV^CR6^ but not MNoV^CW3^ infection. This led us to identify intestinal tuft cells, the only epithelial cell type that expresses CD300lf as a major target cell of MNoV^CR6^ (*29*). Tuft cells are critical regulators of epithelial response to intestinal helminth infection as they are the primary producers of IL-25 (*39, 40*). However, it remains unknown whether tuft cells are essential for infection. In contrast, MNoV^CW3^ does not require tuft cells to establish infection (*38*). MNV-1 or MNoV^CW3^ infection has been reported in dendritic cells, macrophages, monocytes, and B cells *in vivo* (*11, 30, 41*). In *Stat1*-deficient mice, MNoV^CW3^ has been observed in rare epithelial cells consistent with tuft cells (*12*). While not all MNoV strains require tuft cells, we recently showed that CD300lf is essential for infection of diverse MNoV strains *in vivo (28)*.

To elucidate the cellular tropism of MNoV *in vivo*, here we developed a *Cd300lf* conditional knockout (*CD300lf*^*F/F*^) mouse to specifically delete CD300lf from putative MNoV target cells. We demonstrate that CD300lf expression on tuft cells is essential for MNoV^CR6^ but not MNoV^CW3^ infection and this tropism depends on the viral NS1 protein. We find that ablation of CD300lf on myelomonocytic (LysM+) cells reduces MNoV^CW3^ infection in immunocompetent mice. We did not observe any reduction in MNoV^CW3^ infection with depletion of CD300lf in epithelial cells, dendritic cells, neutrophils, or B cells. Interestingly, the anti-viral effects of CD300lf disruption on LysM+ cells were not observed in *Stat1*-deficient mice demonstrating that CD300lf and the innate immune system coordinately regulate MNoV tropism. This suggests CD300lf expression on tuft cells is required for MNoV^CR6^ infection, multiple cell types are sufficient for MNoV^CW3^ infection, and innate immunity regulates MNoV^CW3^ tropism.

## RESULTS

### CD300lf expression on epithelial cells is required for MNoV^CR6^ but not MNoV^CW3^ infection

To elucidate the cellular tropism of MNoV, we generated a *Cd300lf* conditional knockout mouse via CRISPR/Cas9. We inserted LoxP sites flanking exon 3 of *Cd300lf* (*CD300lf*^*F/F*^) which encodes amino acids 129-158 in the ectodomain of CD300lf (Fig 1A). We first crossed this mouse to an epithelial cell specific Cre mouse (*Vil1Cre*) to generate mice with epithelial cells deficient in CD300lf. We validated the efficiency of CD300lf deletion in intestinal epithelial cells by flow cytometry (Fig 1B). CD300lf+ events were significantly enriched in *CD300lf*^*F/F*^ *Vil1Cre-* mice compared to *CD300lf*^*F/F*^ *Vil1Cre+* littermate control mice (Fig 1B). We then challenged these mice with 10^6^ PFU MNoV^CR6^ and MNoV^CW3^ perorally (PO), harvested mesenteric lymph node (MLN), spleen, ileum, and colon and assessed viral genome copies via quantitative real-time PCR (qPCR). At 7 days post-infection (dpi), MNoV^CR6^ genome copies were significantly reduced in the MLN, ileum, and colon of *CD300lf*^*F/F*^ *Vil1Cre+* mice compared to *Cre-* controls (Fig 1C, E-F). MNoV^CR6^ was not detected in the spleens of either *Cre-* or *Cre+* mice consistent with wild type mice studies prior. MNoV^CR6^ was also undetectable in the feces of *Vil1Cre+* mice unlike *Cre-* controls (Fig 1G). Interestingly, MNoV^CR6^ genomes were detected in the MLN, ileum, colon, and feces in a single *CD300lf*^*F/F*^ *Vil1Cre+* mouse (Fig 1C, E-G) whether this reflects emergence of a novel viral variant is unclear.

**Figure 1.**
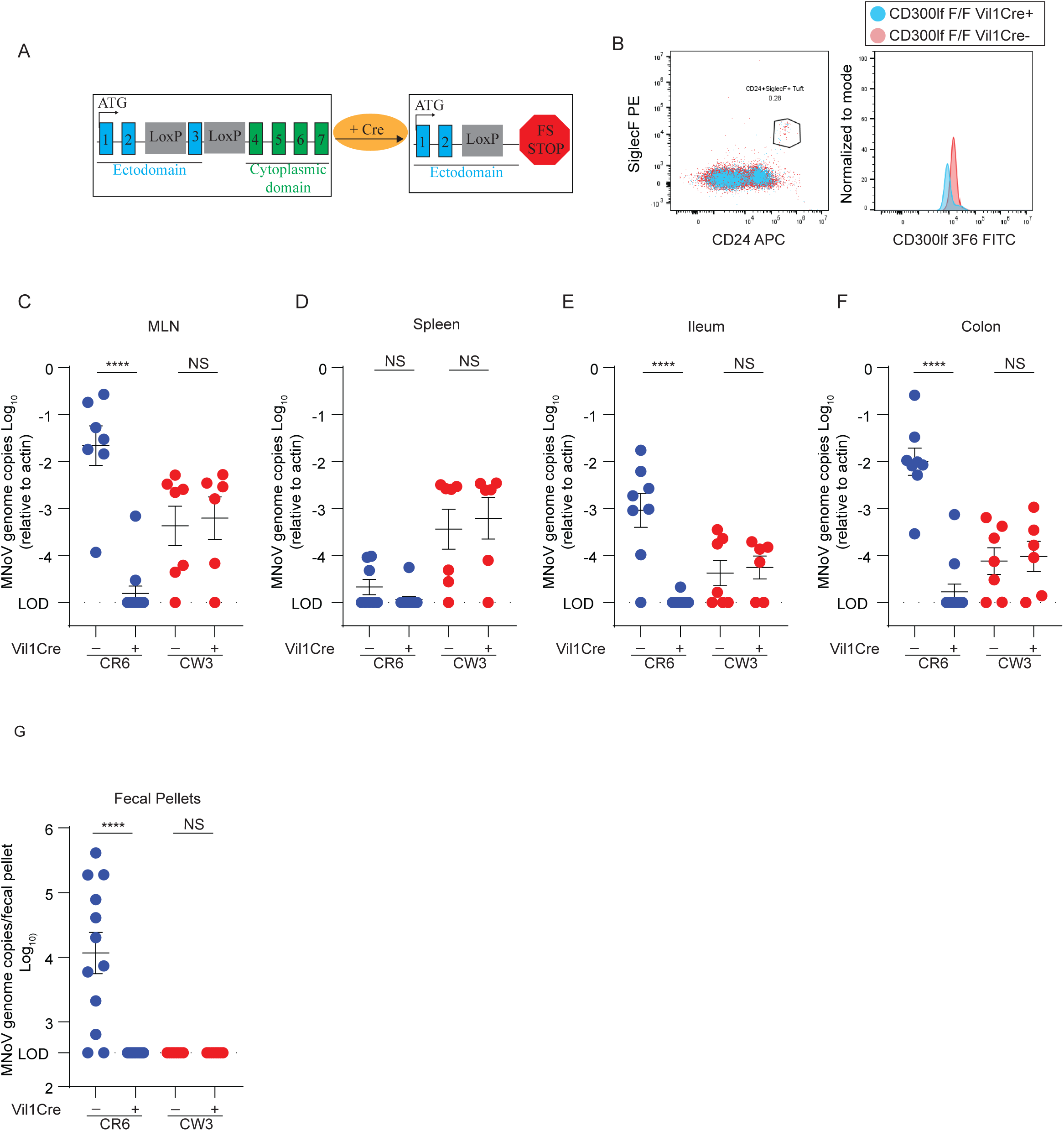
MNoV^CR6^, but not MNoV^CW3^, infection requires CD300lf-expressing epithelial cells. A) Schematic depicting the *Cd300lf* gene locus used to cross with specific cell lineage mouse strains. B) CD300lf expression is ablated on *Vil1Cre+* mice as observed via FACS. (C to F) *CD300lf* ^*F/F*^ *Vil1Cre* mice were perorally infected with 10^6^ PFU of MNoV^CR6^ or MNoV^CW3^ and sacrificed at 7 days post-infection (dpi). Tissue titers for MLN (C), spleen (D), ileum (E), or Colon (F), were analyzed via qPCR for MNoV genome copies and normalized to actin. Fecal pellets (G) collected at 7 dpi were analyzed via qPCR for MNoV genome copies. Mouse experiments were performed using littermate controls with at least two independent repeats analyzed via Mann-Whitney test. Statistical significance annotated as: ***, p < 0.001, NS = not significant.

In contrast to MNoV^CR6^, MNoV^CW3^ viral genome was detected at similar levels in the MLN and spleen of *CD300lf*^*F/F*^ *Vil1Cre+* and *Vil1Cre-* mice (Fig 1C-D). MNoV^CW3^ viral genomes were also detected at similar levels in the ileum and colon of *CD300lf*^*F/F*^ *Vil1Cre-* and *Cre+* mice (Fig 1E-F). MNoV^CW3^ did not robustly shed in the feces of either *Cre-* or *Cre+* mice consistent with prior findings (Fig 1G) (*25, 29*). Together, these results suggest that productive infection of IECs is not required for MNoV^CW3^ infection. Together this suggests that CD300lf expression on epithelial cells is essential for fecal-oral transmission of MNoV^CR6^ but not MNoV^CW3^ (*29*).

### CD300lf expression on tuft cells is required for MNoV^CR6^ and MNoV^CW3-NS1-CR6^ but not MNoV^CW3^

Recently, we showed that MNoV^CR6^ infects rare IECs called tuft cells (*29*). To test whether tuft cells were essential for MNoV infection, we crossed *CD300lf*^*F/F*^ mice to a double cortin-like kinase 1 (Dclk1) Cre – specific to tuft cells - to ablate CD300lf infection on tuft cells (*42*). We then infected *CD300lf*^*F/F*^ *Dclk1Cre-* and *Cre+* mice with 10^6^ PFU PO of MNoV^CR6^, MNoV^CW3^ and chimeric MNoV^CW3^ expressing the NS1 of MNoV^CR6^ (MNoV^CW3-NS1-CR6^). Mice were then sacrificed at 7 dpi for tissue analysis. Consistent with the *CD300lf*^*F/F*^ *Vil1Cre* mice, MNoV^CR6^ was able to infect *CD300lf*^*F/F*^ *Dclk1Cre-* mice but failed to infect *CD300lf*^*F/F*^ *Dclk1Cre+* animals confirming that tuft cells are the exclusive site for MNoV^CR6^ infecton (Fig 2A-D). Both *CD300lf*^*F/F*^ *Dclk1Cre-* and *Cre+* were susceptible to MNoV^CW3^ suggesting that CD300lf expression on tuft cells is not essential for MNoV^CW3^ pathogenesis (Fig 2A-D). Interestingly, CD300lf disruption on tuft cells reduced MNoV^CW3-NS1-CR6^ viral loads in the intestinal tissue but did not have a significant effect on MNoV^CW3-NS1-CR6^ infection in systemic tissues (Fig 2A-D). This is consistent with the NS1 of MNoV^CR6^ enabling tuft cell tropism and the VP1 of MNoV^CW3^ promoting systemic infection (*12, 25, 26, 31*).

**Figure 2.**
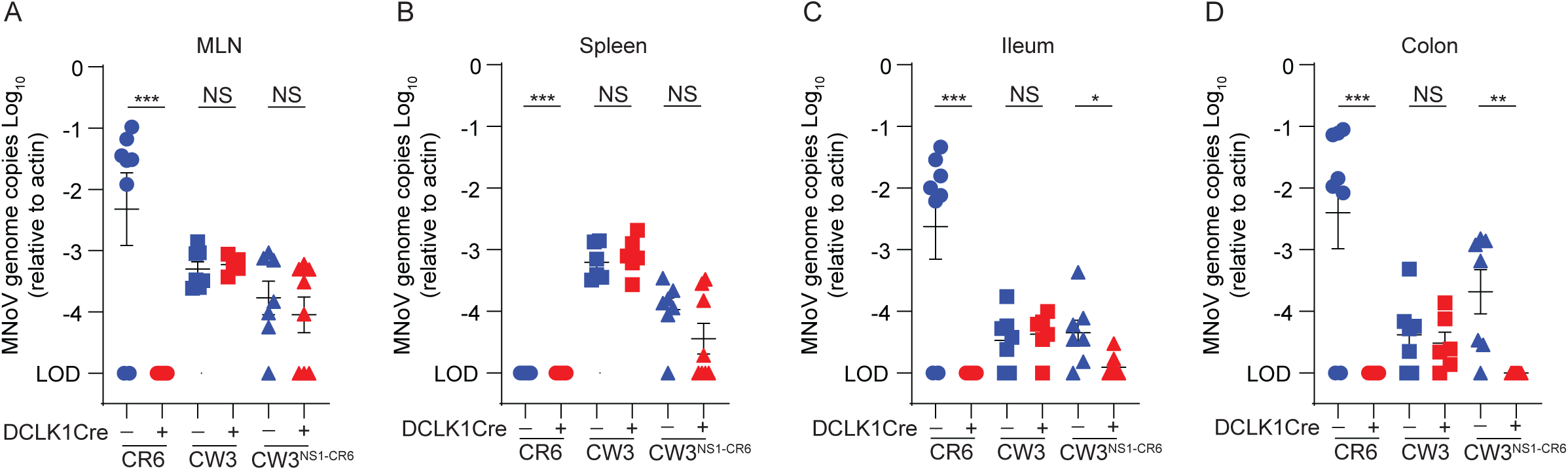
CD300lf expression on tuft cells is required for MNoV^CR6^ but not MNoV^CW3^ or MNoV^CW3-NS1-CR6^ infection. *CD300lf* ^*F/F*^*DCLK1Cre-* and *Cre+* mice were infected with 10^6^ PFU PO of MNoV^CR6^, MNoV^CW3^, and MNoV^CW3-NS1-CR6^ and sacrificed at 7 days post-infection to assess viral titers in the MLN (A), spleen (B), ileum (C), and colon (D) by qPCR. MNoV genome copies were then normalized to actin. At least 2-3 independent repeats were performed using littermate controls. Statistical analysis was performed using Mann-Whitney test. Significance is annotated as: ***, p < 0.001; **, p < 0.01; *, p < 0.05; NS = not significant.

### B cells and dendritic cells are not required for MNoV^CW3^ infection

We next evaluated the role of CD300lf-expressing hematopoietic cell types previously implicated in MNoV^CW3^ infection. B cells have been suggested as a major target cell of both HuNoV and non-persistent MNV both *in vivo* and *in vitro* (*11, 41, 43*). To test whether B cells were essential for MNoV^CW3^ infection we generated a *CD300lf*^*F/F*^ *CD19 Cre* line. CD19 is only transcribed in cells of B lineage and is expressed throughout B cell development and differentiation (*44*). 10^6^ PFU of MNoV^CW3^ was administered PO to these mice and tissue samples were harvested at 7 dpi for qPCR. There was no significant difference in viral genomes between *CD300lf*^*F/F*^ *CD19 Cre+* and *Cre-* littermates indicating CD300lf on B cells is not essential for productive MNoV^CW3^ infection (Fig 3A-D).

**Figure 3.**
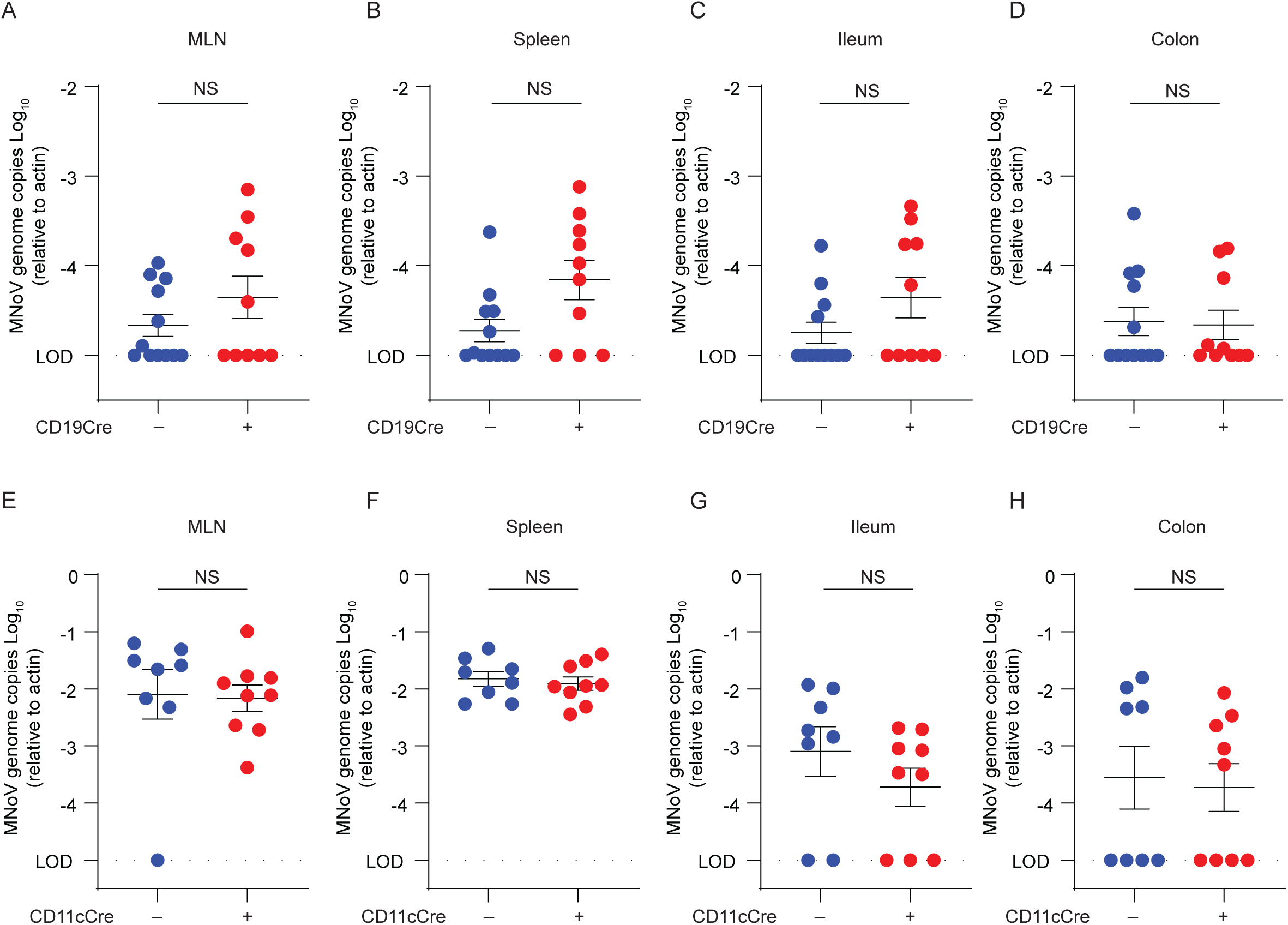
CD300lf expression on B cells (*CD19 Cre+)* and dendritic cells (*CD11c Cre+)* is not essential for MNoV^CW3^ infection *in vivo*. (A-H) *CD300lf* ^*F/F*^ *CD19Cre* and *CD300lf* ^*F/F*^ *CD11cCre* mice were perorally infected with 10^6^ PFU of MNoV^CW3^ and sacrificed at 7 days post infection. MNoV titers for the MLN (A), spleen (B), ileum (C), colon (D) were analyzed via qPCR for MNoV genome copies and normalized to actin. Mouse experiments were performed using littermate controls with four independent repeats analyzed via Mann-Whitney test. Statistical significance annotated as: NS = not significant.

As dendritic cells have also been implicated as a target cell of non-persistent MNoV^CW3^ (*41, 43, 44*), we generated *CD300lf*^*F/F*^ *CD11cCre* mice to test the role of dendritic cell infection in MNoV^CW3^ pathogenesis. CD11c, also known as integrin αX, is a widely used marker for dendritic cells (*45*). Similar to CD19-Cre, there was no significant difference in MNoV^CW3^ viral genome copies in the MLN, spleen, ileum, or colon (Fig 3E-H), suggesting CD300lf expression on dendritic cells is not essential for productive MNoV^CW3^ infection *in vivo*.

### LysM positive cells contribute to MNoV^CW3^ infection

To ascertain whether myelomonocytic cells were essential for MNoV^CW3^ infection, we generated mice with myeloid lineage cells deficient in CD300lf (*CD300lf*^*F/F*^ *LysM Cre)*. LysM is a lysozyme that is widely produced by immune cells and serves as a marker for myelomonocytic cells (*46, 47*). CD300lf expression on WT bone marrow macrophages (BMDMs) is below the limit of detection by flow cytometry (*27*). Therefore, we validated the activity of the Cre recombinase by harvesting BMDMs from both *CD300lf*^*F/F*^ *LysM Cre-* and *Cre+* mice and infecting these cells with MNoV^CW3^ at a MOI of 0.05. We quantified viral replication by plaque assay at 1- and 24-hours post-infection (hpi). BMDMs from *CD300lf*^*F/F*^ *LysM Cre+* mice have reduced infectious virus as compared to BMDMs from *LysM Cre-* littermates at 24 hpi consistent with efficient CD300lf disruption in macrophages (Fig 4A). *CD300lf*^*F/F*^ *LysM Cre-* and *Cre+* mice were then challenged 10^6^ PFU MNoV^CW3^ PO and tissues were harvested at 7 dpi for qPCR. There was a significant reduction in viral genomes in spleen and ileum in *CD300lf*^*F/F*^ *LysM Cre+* mice (Fig. 4B-E), suggesting that MNoV^CW3^ infection is in part supported by LysM positive cells *in vivo*. Interestingly no significant difference in viral genomes was detected between *CD300lf*^*F/F*^ *LysMCre-* and *Cre+* littermates in the MLN and colon.

**Figure 4.**
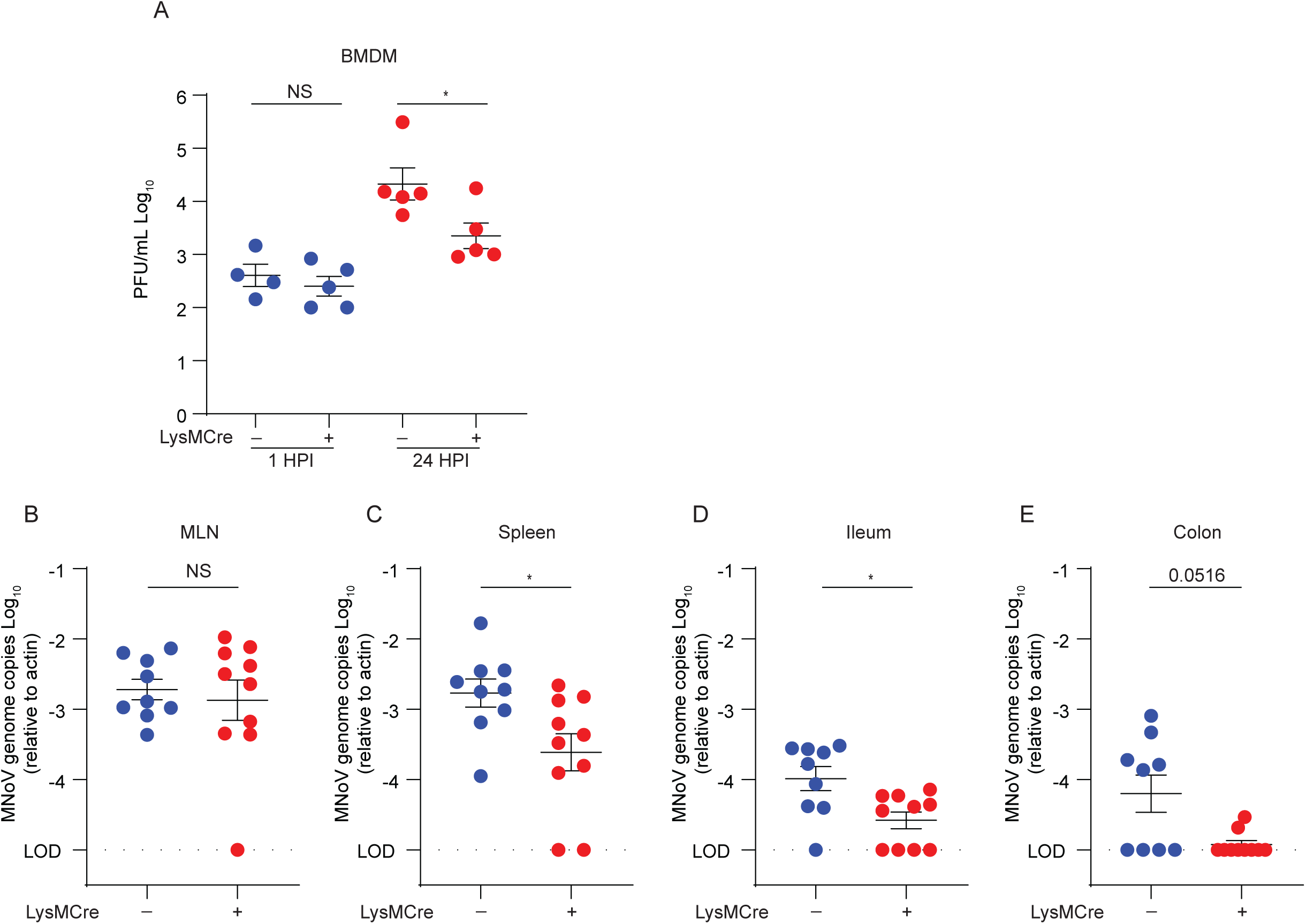
LysM+ cells contribute to productive MNoV^CW3^ infection in a CD300lf-dependent manner. (A) Bone marrow-derived macrophages from *Cd300lf*^*-/-*^ and *CD300lf* ^*F/F*^ *LysMCre* mice were infected with MNoV^CW3^ at a MOI of 0.05 for 1 or 24 hours post-infection. Infectious virus was quantified by plaque assay. (B to E) *CD300lf* ^*F/F*^ *LysMCre* mice were perorally infected with 10^6^ PFU of MNoV^CW3^ and sacrificed at 7 days post-infection. Tissue titers for the MLN (B), spleen (C), ileum (D), colon (E) were analyzed via qPCR for MNoV genome copies and normalized to actin. Mouse experiments were performed using littermate controls with four independent repeats analyzed via Mann-Whitney test. Data in A was analyzed via unpaired students t-test. Statistical significance annotated as: *, p < 0.05; **, p < 0.01, ****, p < 0.0001 NS = not significant.

### CD300lf expression on neutrophils is not essential for MNoV^CW3^ infection

LysM is expressed on macrophages and neutrophils, which both express high levels of CD300lf (48). To determine whether neutrophils are infected by MNoV^CW3^, we crossed the *CD300lf*^*F/F*^ to a myeloid-related protein 8 (Mrp8) mouse to generate mice with CD300lf deficient neutrophils (*CD300lf*^*F/F*^ *Mrp8 Cre+*). Blood samples were collected from these mice and CD300lf disruption efficiency was assessed by flow cytometry (Fig 5A). Samples harvested from *CD300lf*^*F/F*^ *Mrp8 Cre+* mice showed a reduction in CD300lf on neutrophils as compared to the *CD300lf*^*F/F*^ *Mrp8 Cre-* mice (Fig 5B-C), confirming activity of the Cre recombinase. These mice were then infected PO with 10^6^ PFU of MNoV^CW3^ and tissues were collected at 7 dpi (Fig 5D-G). Viral genome copies in the MLN, spleen, ileum and colon were equivalent between *CD300lf*^*F/F*^ *Mrp8 Cre-* and *Cre+* mice (Fig 5D-G). These results suggest that neutrophils are not required for MNoV^CW3^ infection *in vivo*.

**Figure 5.**
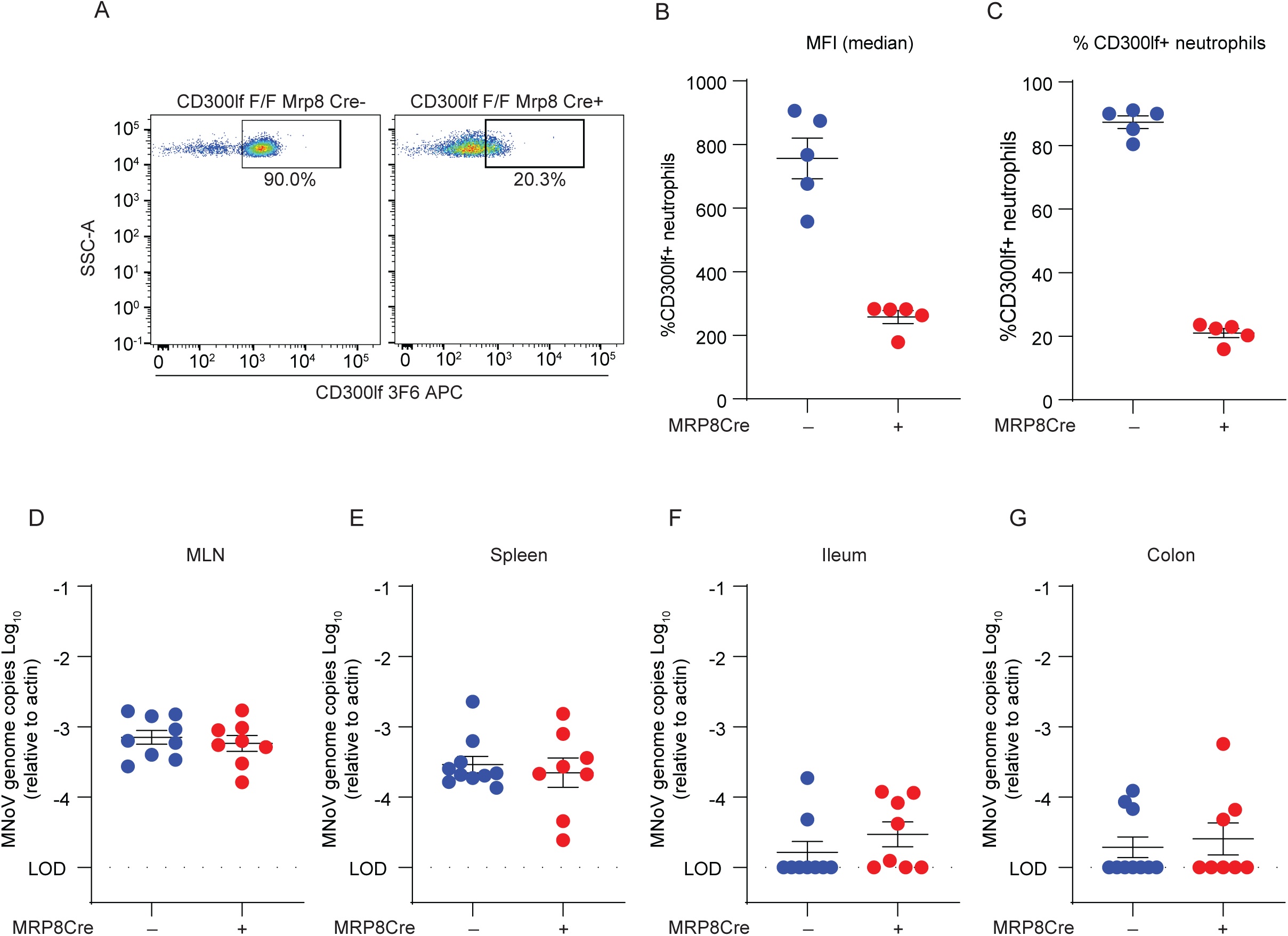
CD300lf depletion on neutrophils does not affect MNoV^CW3^ infection *in vivo*. (A) FACS plot and quantification of CD300lf expression levels on neutrophils harvested from *CD300lf* ^*F/F*^ *MRP8 Cre* mice. Each dot represents one mouse. (B-E) *CD300lf* ^*F/F*^ *MRP8Cre* mice were infected with 10^6^ PFU PO of MNoV^CW3^ and sacrificed at 7DPI. MNoV^CW3^ genome copies from MLN (B), spleen (C), ileum (D), colon (E) were analyzed by qPCR and normalized to actin. Data in A was analyzed via Mann-Whitney test and is representative of at least two independent experiments. Mouse experiments were performed using littermate controls with two independent repeats analyzed via Mann-Whitney test. Statistical significance annotated as: **, *p* < 0.01; NS = not significant.

### MNoV^CW3^ infection of LysM-positive cells is not essential for lethality in innate immune deficient mice

Mice deficient in type I interferon signaling or transcription factor STAT1 are highly susceptible to MNoV^CW3^ infection resulting in lethality (*23, 48*). Given that LysM+ cells contribute to MNoV^CW3^ infection, we asked whether disruption of CD300lf on LysM+ cells would confer protection against MNoV^CW3^ on a *Stat1*^*-/-*^ background. We challenged *CD300lf*^*F/F*^ *LysMCre-Stat1*^*-/-*^or *Cre+ Stat1*^*-/-*^ mice 10^4^ or 10^6^ PFU MNoV^CW3^ and assessed survival. No significant differences in lethality were observed between these lines at either viral challenge dose (Fig 6A-B). Next, we harvested tissues for viral RNA quantification at 3 dpi, prior to the onset of lethality. Interestingly, viral genomes were equivalent between the *CD300lf*^*F/F*^ *LysM Cre-Stat*^*-/-*^ and *CD300lf*^*F/F*^ *LysM Cre+ Stat*^*-/-*^ mice across tissues (Fig 6C-F). These results indicate that in the absence of interferons, MNoV^CW3^ infection is not dependent on the expression of CD300lf on LysM positive cells.

**Figure 6.**
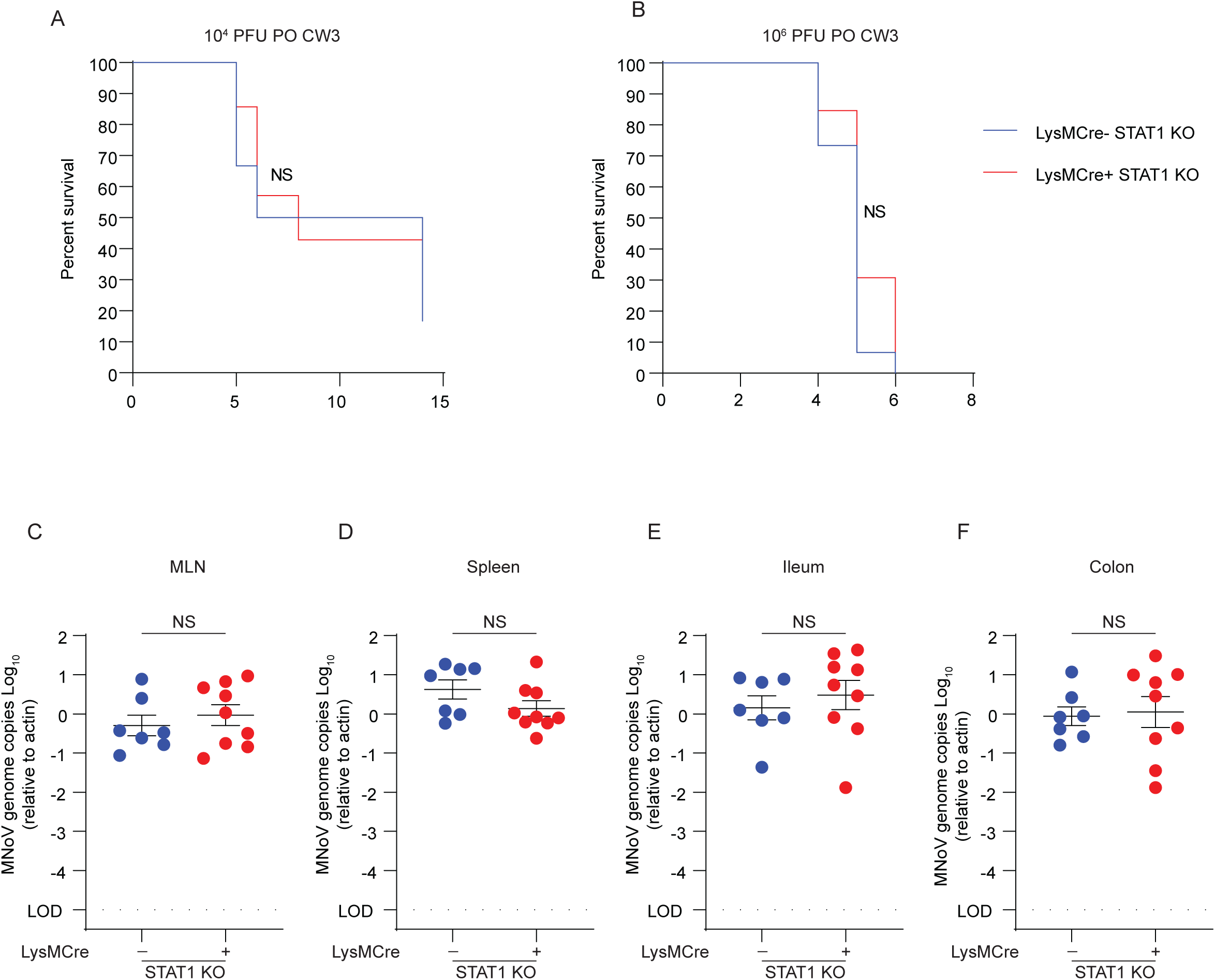
Ablation of CD300lf expression on LysM^+^ cells does not provide protection from lethal MNoV^CW3^ infection in *Stat1*^*-/-*^ mice. (A-B) *CD300lf* ^*F/F*^ *LysMCre* x *Stat1*^-/-^mice were infected with either 10^4^ (A), or 10^6^ (B) PFU of MNoV^CW3^ and observed for lethality for 14 days post-infection (dpi) or 7 dpi respectively. (C-F) *CD300lf* ^*F/F*^ *LysMCre* mice were infected with 10^6^ PFU PO of MNoV^CW3^ and sacrificed at 3 dpi. Tissue titers for the MLN (C), spleen (D), ileum (E), colon (F) were analyzed via qPCR for MNoV genome copies and normalized to actin. Mouse experiments were performed using littermate controls with pooled data from two-three independent repeats and analyzed via Mann-Whitney test. Kaplan-Meier curves were generated for survival experiments. Statistical significance annotated as: NS = not significant.

## Discussion

We previously identified CD300lf as the primary physiologic receptor for MNoV (*27, 29*). CD300lf is necessary and sufficient for infection of diverse MNoV strains *in vivo* which we leverage to interrogate MNoV cell tropism (*27, 29*). Here, we introduce a novel conditional knockout mouse model that serves as a valuable tool for studying tissue specific CD300lf expression. We previously demonstrated that persistent MNoV^CR6^ infects a rare population of IECs called tuft cells. However, it was unknown whether tuft cell infection is essential *in vivo* and infection as other putative target cells may support MNoV^CR6^ replication. Here, we show that that MNoV^CR6^ infection can be ablated by conditionally deleting CD300lf on IECs broadly and tuft cells more specifically. This demonstrates that CD300lf expression on tuft cells is essential for per oral infection of MNoV^CR6^. In contrast to MNoV^CR6^, MNoV^CW3^ was unaffected by conditional ablation of CD300lf on both IECs and tuft cells-specifically. Using a chimeric virus (MNoV^CW3-NS1-CR6^), we demonstrate that the NS1 of CR6 permits tuft cell tropism, which requires tuft cell-specific CD300lf expression for intestinal tissue infection (*12, 25, 29, 31*). Interestingly, MNoV^CW3-NS1-CR6^ maintained the ability to infect systemic tissues, suggesting additional determinants of cell and tissue tropism for MNoV. These data ultimately support different tropism patterns across MNoV strains.

Previous reports identified dendritic cells, B cells, macrophages, monocytes, and neutrophils as targets of MNoV^CW3^(*43, 49, 50*). We therefore sought to investigate the relative contribution of these cell types to MNoV^CW3^ infection by crossing our *CD300lf*^*F/F*^ to cell specific Cre recombinases to generate mouse lines with CD300lf deleted on specific target cells. We demonstrate that CD300lf-expression on LysM+ cells (i.e.macrophages and monocytes) significantly contribute to MNoV^CW3^ infection, whereas we observed no impact of CD300lf disruption on dendritic cells, B cells, and neutrophils. Our finding that LysM+ cells are important for MNoV^CW3^ infection is consistent with a recent report demonstrating that MNoV^CW3^ requires susceptible myeloid cells and depends on cell lysis to induce chemotaxis and inflammatory responses for myeloid cell recruitment to the site of infection (*50*). Our observation of only a partial reduction in MNoV^CW3^ viral load in mice in which CD300lf is ablated from LysM+ cells supports that a single target cell type may not be essential for MNoV^CW3^ infection, as it is for MNoV^CR6^. Instead, multiple CD300lf-expressing cell types likely contribute.

MNoV tropism is governed both by virus-receptor and virus-immune system interactions. Specifically, disruption of type I IFN signaling through IFNAR or STAT1 deletion results in lethal MNoV^CW3^ infection and systemic spread of MNoV^CR6^ likely due to expanded cell tropism (*23, 26*). Consistent with this, we observed no reduction of MNoV^CW3^ viral load in *Stat1*^*-/-*^ mice with conditional ablation of CD300lf on LysM+ cells demonstrating that STAT1 partially restricts MNoV^CW3^ to myelomonocytic cells and suggests an expanded tropism in mice lacking innate immunity. The mechanisms of this restriction represent an important area of future investigation.

In addition to its role in MNoV infection, CD300lf has been implicated in diverse disease states including multiple sclerosis, inflammatory bowel disease, and depression (*33, 36-38, 51, 52*). Elucidating the cell-type specific role of CD300lf in these disease contexts may provide critical insight into mechanisms of pathogenesis. The mouse model described herein thus provides an important tool for studying CD300lf-expression in various contexts.

HuNoV cell tropism remains incompletely understood. Diverse hematopoietic and epithelial cell types including enteroendocrine cells have been implicated in HNoV infection *in vivo* in humans, non-native hosts, and *in vitro* (*7, 9-11, 18*). Elucidating the host and viral determinants governing HNoV tropism may reveal insight into HNoV transmission, pathogenesis, and persistence. MNoV provides a useful tool to reveal molecular interactions at the viral and host levels that may inform studies of HNoV.

## Acknowledgments

We would like to acknowledge Skip Virgin and Darren Kreamalmayer for their generous resources. **Funding:** This work was supported by NIH grants K08 A1128043 (CBW) and R01 AI127552 (MTB) and R01 AI139314 (MTB). **Author contribution: Vincent R. Graziano;** conceptualization, formal analysis, investigation, writing-original draft, visualization, **Mia Madel Alfajaro;** validation, formal analysis, investigation, writing-original draft, visualization, **Cameron O. Schmitz;** formal analysis, investigation, **Renata B Filler;** investigation, **Madison S. Strine;** formal analysis, investigation, **Jin Wei;** investigation, **Leon L. Hsieh;** investigation, **Megan T. Baldridge;** conceptualization, supervision, **Timothy J. Nice;** conceptualization, supervision, **Sanghyun Lee;** conceptualization, supervision, **Robert C Orchard;** conceptualization, methodology, supervision, resources; **Craig B Wilen;** conceptualization, methodology, supervision, investigation, resources, funding acquisition, writing-original draft. All authors reviewed and edited the manuscript.

## Materials and Methods

### Mouse Strains

Cd300lf conditional knockout mouse were generated by introducing LoxP sites flanking exon 3 of Cd300lf. The 5’ and 3’ LoxP sites were introduced by CRISPR/Cas9 mediated homology-directed repair. The following single guide RNAs (sgRNAs) were used: GTGTTGTGGCCTAACTTGCANGG and TCAAGTTCCCTGTCTCTTGGGGG to insert donor sequence GAATTCATAACTTCGTATAATGTATGCTATACGAAGTTAT at position Chr11: 115,122,845-115,122,846 and donor sequence GTGCTCATTAATGATGTTCTCTTTGAGAGTCCTTCTAGAG at Chr 11: 115,125,303-115,125,304. Mice were genotyped by qPCR for the 5’ LoxP site at Transnetyx, Inc.

### Ethics Statement

Animal use and care was approved in agreement with the Yale Animal Resource Center and Institution Animal Care and Use Committee (#2018-20198) according to the standards set by the Animal Welfare Act.

### Viral Stocks

Molecular clones of MNoV^CW3^ (Gen Bank accession EF014462.1) and MNoV^CR6^ (Gen Bank accession JQ237823) we used to generate a working stock of infectious virus. To create stocks, plasmids containing infectious molecular clones were transfected into 293T cells (ATCC) to generate a P0 stock as described previously(*27*),(*26*). Then, the P0 stock was used to infect susceptible BV2 cells (gifted from H. W. Virgin) for generation of the P1 stock. Generation of the P2 stock was performed by inoculating BV2 cells with the P1 stock for 36 hours followed by freezing the infected cultures at −80°C. Upon thawing the infected BV2 cultures, cellular debris was pelleted at 1200g for 5 minutes (min) and then filtered through a 0.22 µm filter and subsequently concentrated via 100,000 MWCO Amicon Ultra Filter. Concentrated viral stocks were aliquoted and stored at −80°C. Further tittering was performed via plaque assay at least three independent times.

### Cell lines culture

BV2 cells were maintained in Dulbecco’s modified eagle media (DMEM; Gibco, Gathersburg, MD) that was supplemented with 10% fetal bovine serum (FBS; VWR, Radnor, PA), 1% HEPES (Gibco), and 1% Penicillin/Streptomycin (Pen/Strep) (Gibco). In order to differentiate progenitors into BMDMs, 10^6^ cells were plated into a 10 cm non-tissue culture treated plate with BMDM media (DMEM, 10% FBS, 10% CMG14 conditioned media, 2 mM L-glutamine, 1% Pen/Strep, and 1% sodium pyruvate), incubated at 37°C for seven days at 5% CO2 (*53*). Differentiation of progenitors into BMDMs was confirmed via flow cytometry by staining for F4/80.

### *In Vitro* MNoV infections

BMDMs from *Cd300lf*^*-/-*^ and *Cd300lf*^*f/f*^ *LysMCre* mice were harvested as described above. After 7 days in differentiation media, 50,000 cells were plated per well of a 96-well plate and infected with MNoV^CW3^ at a MOI of 0.05. Infected plates were frozen at −80°C at 1 and 24 hpi for plaque assay.

### *In Vivo* MNoV Infections

Mouse infections were performed by inoculating with 25 µL of 10^6^ PFU MNoV^CW3^ or MNoV^CR6^ diluted in DMEM supplemented with 10% FBS. At 7 days post infection, mice were sacrificed and fecal samples as well as tissues were stored at −80°C until processing.

### Viral quantification by plaque assay

Each well of a 6-well plate was seeded with 2 ⨯10^6^ BV2 cells in DMEM with 10% FBS, 1% Pen/Strep, 1% HEPES and then incubated overnight at 37°C and 5% CO2. After 24 hours, the BV2 cells were checked for confluency and BMDM-infected plates were thawed followed by 6 serial dilutions. The media was removed from the BV2 cells and one diluted inoculum was added to one well of the 6-well plate and rocked gently for 1 hour. After rocking, the inoculum was removed, and 2 mL overlay media was applied (MEM with 10% FBS containing 1% methylcellulose, 1% HEPES, 1% GlutaMAX (Gibco), and 1% Pen/Strep) followed by a 48 hours incubation. Following incubation, overlay media was aspirated off and each well was stained with 1 mL crystal violet (20% ethanol, and 0.2% crystal violet) on a plate rocker for 30 min as previously described (*41*).

### Quantitative PCR

Viral genome copies in tissues and fecal pellets were previously described (*27, 54*). Briefly, viral RNA extraction from tissues was performed using TRIzol (Life Technologies, Carlsbad, CA) and purified with a Direct-Zol RNA MiniPrep Plus kit according to manufacturer’s protocol (Zymo Research, Irvine, CA). Following purification, a two-step cDNA synthesis was performed using 5 µL of RNA, random hexamers and ImProm-II Reverse Transcriptase (Promega) was performed. qPCR analysis on standard curves and samples was performed in duplicate using MNoV specific oligonucleotides: Forward primer: 5’ CACGCCACCGATCTGTTCTG 3’; Reverse primer: 5’ GCGCTGCGCCATCACTC 3’; Probe: 5’ 6FAM-CGCTTTGGAACAATG-MGBNFQ 3’. The limit of detection for qPCR analysis was 10 MNoV genome copies/1 µL. MNoV genome copies were normalized to the expression of housekeeping gene β-actin detected using murine actin oligonucleotides: Forward primer: 5’ GCTCCTTCGTTGCCGTCCA 3’; Reverse primer: 5’ TTGCACATGCCGGAGCCGTT 3’; and Probe: 5’ 6-JOEN-CACCAGTTC /ZEN/ GCCATGGATGACGA-IABKFQ 3’. The limit of detection for β-actin was 100 copies/1 µL. Undetectable genome copies were set at 0.0001 copies relative to actin.

### Flow Cytometry

Mice were euthanized and sacrificed in compliance with IACUC protocol. Colonic tissue was harvested, opened longitudinally, and washed three times in ice cold 1X DPBS. To dissociate cells, colonic tissue was finely chopped with a razor blade and suspended in stripping buffer (1X DPBS, 5% FBS, 0.1% Pen/Strep, 5 mM EDTA, and 0.5 M DTT). Cells were incubated at 37°C for 15 min and gently agitated at 200 rpm. Dissociated cells were filtered sequentially through 100 μm then 40 μm filters. Cellular filtrate was pelleted by centrifugation (1500 rpm for 5 min at 4°C) and washed once with FACS buffer (1X DPBS, 2 mM EDTA, 2.5% FBS). Cells were then stained for viability with Zombie Aqua (BioLegend) diluted 1:500 in 1X DPBS on ice for 10 min. Samples were centrifuged at 1200xg for 2 min and supernatant was removed. CD300lf primary antibody 3F6 (Armenian Hamster, Genentech) was added at a 1:800 dilution for 20 min at room temperature. Cells were then washed with FACS buffer. To identify epithelial cells expressing CD300lf, cells were stained with the following antibodies diluted in FACS buffer: AlexaFluor 488 Goat anti-Armenian Hamster (1:500, Jackson Laborotory, 127545160, lot 128099 in 50% glycerol), EpCAM (1:200, BioLegend), SiglecF (1:200, BioLegend) and CD24 (1:200, BioLegend), and CD45 (1:200, BioLegend) on ice for 20 min. Cells were washed with FACS buffer, resuspended in 4% paraformaldehyde, and filtered. Samples were analyzed on a CytoFLEX S (Beckman Coulter).

### Statistical Analysis

Statistical analysis was performed in Prism GraphPad version 8 (San Diego, CA). Error bars show the standard error of the mean unless indicated otherwise. For all non-normally distributed data, Mann-Whitney tests were performed whereas normally distributed data was analyzed with an unpaired Students T-test. Kaplan-Meier curves were used to analyze survival data. P-values < 0.05 were considered significant (p < 0.05, *; p < 0.01, **; p < 0.001, ***; p < 0.0001, ****)

### Data Availability

All relevant data are contained within the manuscript. CD300lf F/F mice are available upon request.

